# Fluorescent tagging of *Plasmodium* circumsporozoite protein allows imaging of sporozoite formation but blocks egress from oocysts

**DOI:** 10.1101/2020.10.22.350330

**Authors:** Mirko Singer, Friedrich Frischknecht

## Abstract

The circumsporozoite protein, CSP is the major surface protein of *Plasmodium* sporozoites, the form of malaria parasites transmitted by mosquitoes. CSP is involved in sporozoite formation within and egress from oocysts, entry into mosquito salivary glands and mammalian liver as well as migration in the skin. Antibodies against CSP can stop infection prior to the first round of parasite replication in the liver. CSP consists of different domains and is proteolytically cleaved prior to hepatocyte invasion. Part of CSP has been developed into a licensed vaccine against malaria. Yet, how CSP facilitates sporozoite formation, oocyst egress and hepatocyte specific invasion is still not fully understood. Here, we generated a series of parasites expressing full-length versions of CSP as fusion proteins with the green fluorescent protein. This enabled the investigation of sporozoite formation in living oocysts and revealed a dominant negative function of some GFP-CSP fusions during sporozoite egress.

## Introduction

Transmission of malaria occurs when *Plasmodium* sporozoites are inoculated into the skin during the probing phase of a mosquito bite, before the mosquito starts to suck up blood. These sporozoites are formed in parasitic oocysts at the midgut wall of mosquitoes, egress from oocysts to access the hemolymph, the circulatory liquid of the mosquito, and enter into salivary glands. Here they await transmission within the salivary cavities and canals (Frischknecht et al., 2017). Within the skin, sporozoites migrate rapidly and enter both blood and lymph vessels (Amino et al., 2006). Those entering the blood stream are transported with the circulation and arrest specifically in the liver to exit the blood stream and enter hepatocytes, where a liver stage develops that gives rise to thousands of red blood cell infecting merozoites (Prudencio et al., 2006; Cerami et al., 1992). Many of these steps depend on the GPI (glycosyl-phosphatidyl-inositol)-anchored major surface protein CSP (circumsporozoite protein). Antibodies against CSP can lead to a block in migration in the skin (Aliprandini et al., 2018) and reduction of liver cell invasion (Kisalu et al., 2018; Murugana et al., 2020). Part of CSP has been used to develop the only licensed malaria vaccine RTS,S AS01, Mosquirix (Casares, Brumeanu & Richie, 2010).

Deletion of *csp* blocks sporozoite formation as *csp(-)* sporozoites fail at an early step of sporozoite budding from the plasma membrane of the sporoblast (Menard et al., 1997). Reducing *csp* expression still allows for sporozoite formation, yet those parasites are misshaped and noninfectious (Thathy et al., 2002). CSP contains a signal peptide followed by an N-terminal domain, a repeat region and an adhesive thrombospondin repeat (TSR) and is C-terminally anchored to GPI. In the rodent malaria model parasite *Plasmodium berghei*, deletion of the N-terminus leads to a drop in salivary gland invasion and infection of cells within the skin, where a live-stage like parasite can directly produce merozoites (Coppi et al., 2011). Deletion of the repeat region does initially not affect sporozoite formation but leads to defects in sporozoite maturation causing sporozoite death prior to oocyst egress (Ferguson et al., 2014). Deletion of both, the N-terminal domain and repeat region leads to a severe defect in sporozoite budding possibly because forming sporozoites fail to separate their plasma membrane from others (Ferguson et al., 2014). Mutations within the region II+ at the 5’end of the TSR of CSP lead to fully formed sporozoites that fail to egress and can also not enter into liver cells (Wang, Fujioka & Nussenzweig, 2005). Deletion of the GPI-anchor or replacement of the GPI-anchor with a transmembrane domain led to a similar strong effect on sporozoite formation as deletion of *csp* (Wang, Fujioka & Nussenzweig, Cell Microbiol, 2005).

To enable real time imaging experiments to better understand CSP function we aimed at generating a functional fusion protein of CSP with the green fluorescent protein GFP. Visualization of parasite formation in real time has been achieved by fluorescent light microscopy through GFP-tagging of the plasma membrane P-type ATPase PfATP4 and the putative sphingomyelin synthetase PfSMS1 for the blood stage and plasma membrane protein PMP1 for all replicating stages (Rottmann et al., 2010; Kono et al., 2016; Burda et al., 2017). Here we generated a series of parasites expressing versions of CSP “internally” fused to GFP for the investigation of CSP-localization during *Plasmodium berghei* sporozoite formation. This showed that CSP could be successfully tagged and localized to the surface when GFP was introduced either between the repeat region and TSR or between the TSR and GPI-anchor. Both GFP-fusion proteins allowed full sporozoite formation yet sporozoite egress from the oocysts was blocked. Intriguingly, introducing GFP after the signal peptide led to the early cleavage of GFP and did not result in a surface localized fusion protein. In addition, expressing GFP with a GPI-anchor allowed sporozoite egress and colonization of salivary glands.

## Results

### Generation of multiple GFP-CSP fusion proteins

We previously tagged the sporozoite protein TRAP (thrombosponin related anonymous protein) successfully after the signal peptide (Kehrer et al., 2016), while similar tagging of the sporozoite protein TRP1 (thrombospondin related protein 1) failed to give a functional fusion protein (Klug and Frischknecht, 2017). Considering the multiple functions of the different CSP domains, we thus selected several locations for insertion of the *gfp* sequence into the *csp* gene (Figure 1A, Figure S1). Owing to different transfection strategies we generated the following five parasite lines: GFP-GPI, parasites expressing a protein consisting of the signal peptide of CSP, GFP and the C-terminal 22 amino acids of CSP corresponding to the GPI-anchor sequence. This line was obtained through insertion at a silent locus in chromosome 12 of *P. berghei* strain ANKA (Singer et al., 2015) (Figure 1B, Figure S2). SP-GFP-CSP_add, parasites expressing a CSP-GFP fusion protein with the GFP placed between the signal peptide (SP) and the N-terminus of CSP. This line, expressed SP-GFP-CSP in addition to the endogenous CSP and was obtained by linear insertion of the *sp-gfp-csp* plasmid into the *csp* locus on chromosome 4 (Figure 1A, B, Figure S2). SP-GFP-CSP_rep, where the endogenous *csp* was replaced by the *sp-gfp-csp* gene. R-GFP-CSP, where the GFP was placed between the repeat region and TSR of CSP and TSR-GFP-CSP; where the GFP was placed between the TSR and the GPI-anchor. Exact positioning of the GFP insertion sites for R-GFP-CSP and TSR-GFP-CSP was assisted by the crystal structure of the TSR domain (Doud et al., 2012). In these lines, the fusion proteins were also expressed in addition to the endogenous CSP as obtained by linear insertion of the respective plasmids into the *csp* locus on chromosome 4 (Figure 1A, B, Figure S2). Note that for SP-GFP-CSP_add, the DNA is inserted such that the resulting modified locus features an endogenous *csp* with a truncated 3’UTR and the fluorescent copy a truncated 5’UTR, while this is inversed for R-GFP-CSP and TSR-GFP-CSP (Figure 1B). For SP-GFP-CSP_rep, the gene is flanked by the complete 5’UTR and 3’UTR.

**Figure 1.**
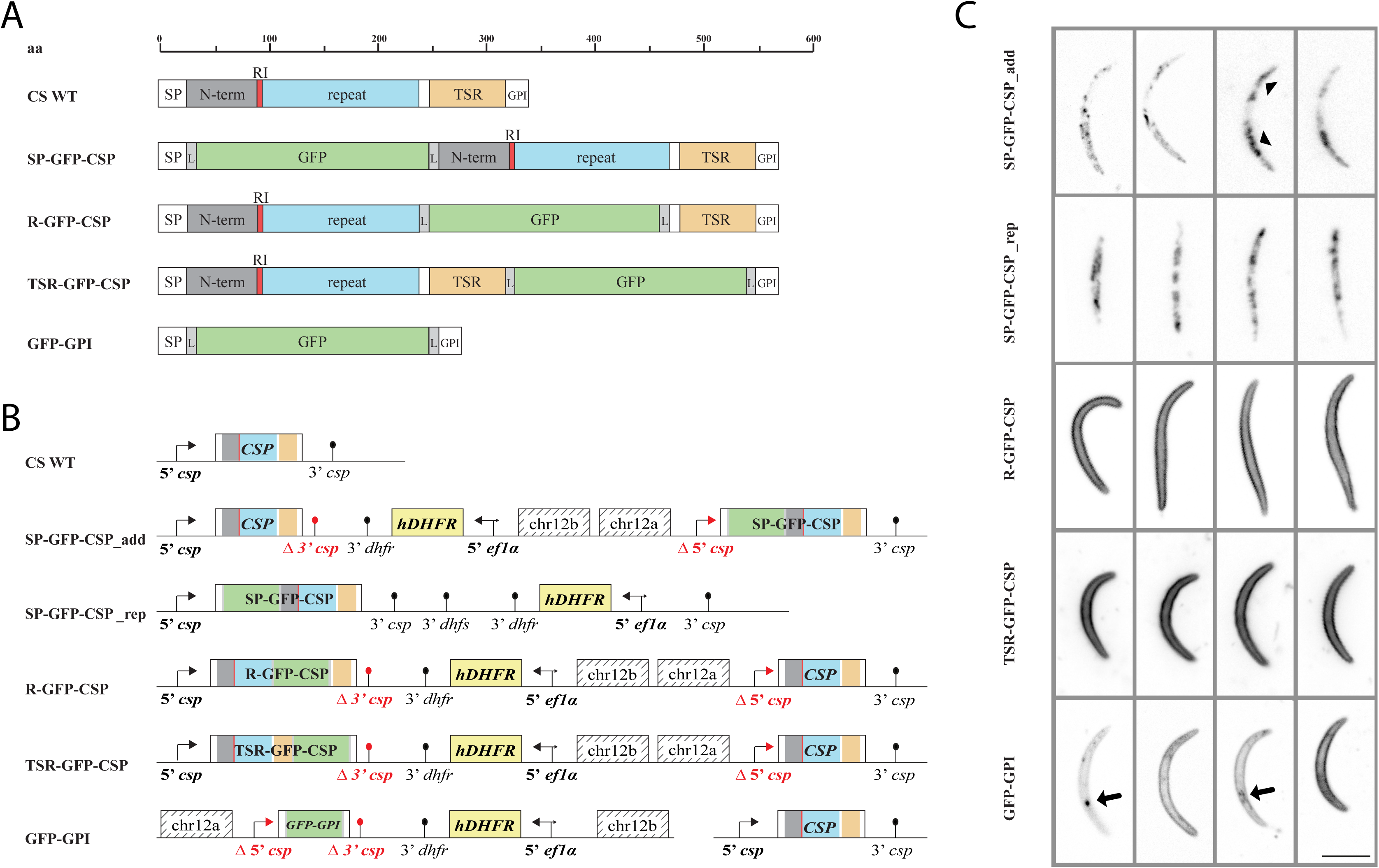
Generation of GFP-CSP fusion lines. A: Cartoon version of the different GFP-CSP fusion proteins showing the respective size of all domains of the circumsporozoite protein (CS) and of all interdomain GFP tags generated. Sizes of respective parts of the proteins are indicated. GFP is directly flanked with a glycine linker (L). B: Gene models of all generated lines. The *csp* locus is indicated for all parasite lines. For GFP-GPI, the *csp* locus is unchanged as the expressed construct is integrated in a silent locus on chromosome 12. Note that for GFP-GPI, both the promoter region Δ5’ csp as well as Δ3’ csp are truncated. In all other lines either the inserted GFP-CSP fusion (SP-GFP-CSP_add) or the endogenous *csp* (R-GFP-CSP and TSR-GFP-CSP) show a shortend 5’UTR (Δ5’ csp). The respective other gene features a shortened 3’UTR (Δ3’ csp) courtesy of the insertion strategy. The SP-GFP-CSP_rep parasite line shows the GFP-CSP fusion with the endogenous 5’ and 3’ UTRs. C: Localization of fluorescence within free GFP-CSP expressing sporozoites. Note the vesicular pattern in the GFP-GPI (arrows) as well as the SP-GFP-CSP (arrowheads) parasite lines. For GFP:GPI, SP-GFP-CSP_add and TSR-GFP-CSP salivary gland sporozoites are shown, for the others midgut sporozoites. The apical end of the sporozoite always points to the bottom. Scale bar: 5 μm.

All lines were readily obtained and generated comparable number of oocysts and midgut sporozoites as wild type parasites, yet some lines showed no sporozoite accumulation in the salivary gland (Table 1). Investigating the GFP-CSP localization revealed that all but the two SP-GFP-CSP lines showed the expected surface localization of the fusion proteins (Figure 1C). Close inspection also revealed that the GPI-GFP expressing parasite often showed additional small vesicular localization in the proximity of the plasma membrane at the estimated nuclear exit site or Golgi localization (Figure 1C, Figure 7).

**Table 1.** Infectivity of the different parasite lines

| Parasite line | MGS | HLS | SGS | HLS/MGS | SGS/MGS | #<sup>1</sup> |
| --- | --- | --- | --- | --- | --- | --- |
| Wild type | 77.000 | 3.400 | 18.000 | 0,03 | 0,2 | 68 (23) |
| GFP-GPI | 46.000 | 500 | 2.500 | 0,01 | 0,05 | 160 (97) |
| SP-GFP-CSP_add | 11.000 | 200 | 4.400 | 0,04 | 0,4 | 154 (119) |
| SP-GFP-CSP_rep | 120.000 | 200 | 10 | 0,005 | 0,0001 | 62 (36) |
| R-GFP-CSP | 29.000 | 100 | 1 | 0,003 | 0,00003 | 97 (97) |
| TSR-GFP-CSP | 21.000 | 200 | 200 | 0,007 | 0,01 | 108 (58) |

MGS: midgut derived sporozoites; HLS: hemolymph derived sporozoites, SGS: salivary gland derived sporozoites; orange and red numbers indicate small and large difference to wild type, respectively. Numbers determined from at least three dissections of at least two infection experiments.
<sup>1</sup>Number of infected mosquitoes analyzed for MGS and SGS as well as for (HLS)

**Figure 2.**
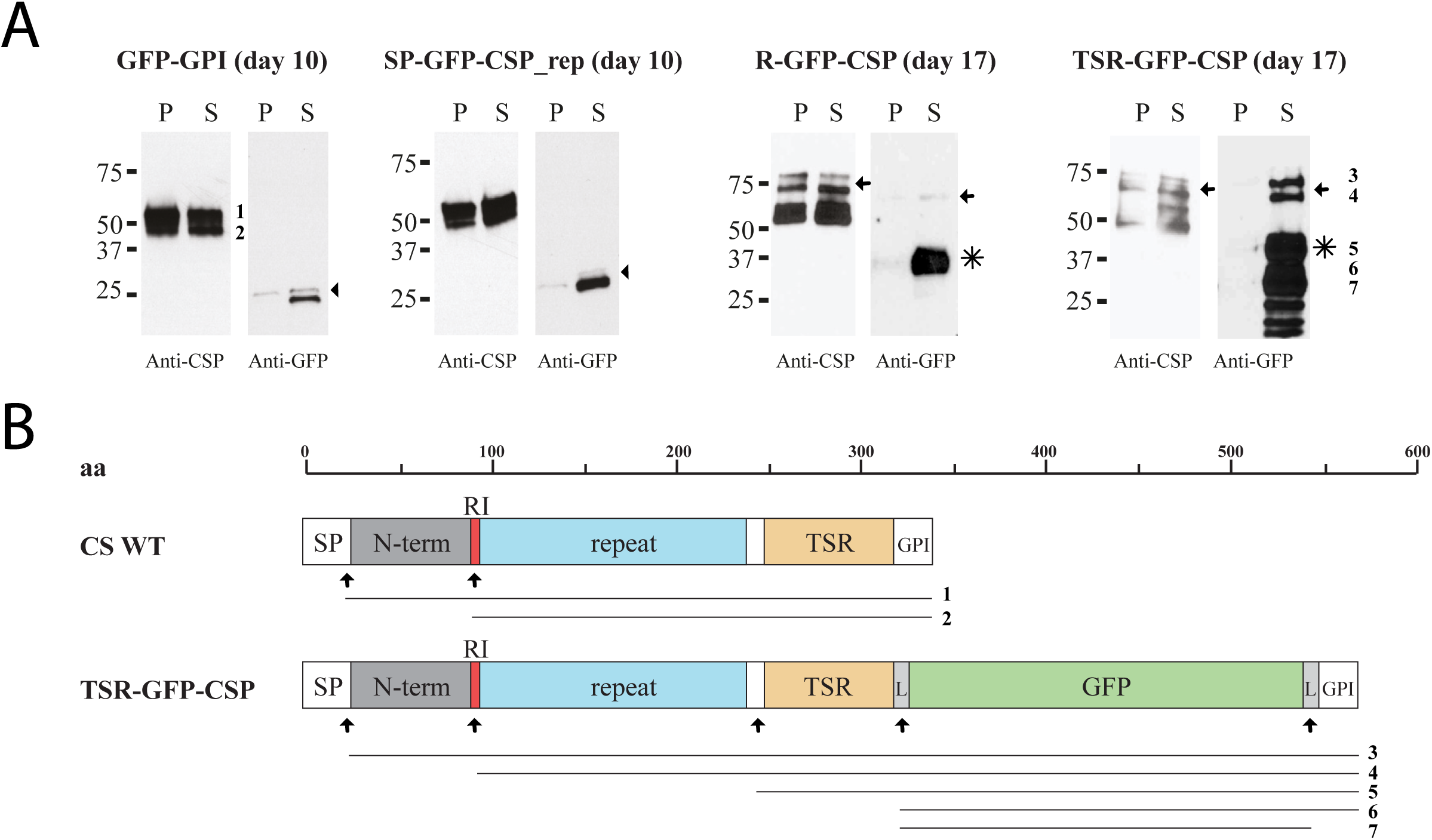
Western blots reveal processing of GFP-CSP fusion proteins. A: Western blots using the 3D11 anti CSP repeats antibody and an anti GFP antibody on the different indicated parasite lines at day 10 or day 17 post infection. Parasites were purified using Accudenz (S) and the pellet (P) was loaded as comparison. Note that in the GFP-GPI and SP-GFP-CSP_rep lines only the typical wild type bands of CSP are revealed by the 3D11 antibody, indicating that no fusion protein is present. In contrast bands corresponding to fusion proteins are readily detectable in the R-GFP-CSP and TSR-GFP-CSP lines (arrows). Note the different degradation products, indicating processing of these GFP-CSP proteins (stars). Arrowheads indicate free or GPI-anchored GFP. Cleavage products shown in B are indicated by small numbers. B: Cartoon illustrating the likely processing events leading to the different bands observed in panel A. Known processing sites after the signal peptide and within region I (RI) are indicated by arrows. Resulting products detectable by anti CSP repeat antibody for CSP WT (1 and 2) and TSR-GFP-CSP (3 and 4, also detectable by anti GFP antibody) are indicated below and also shown in A. Processing/degradation near the linker regions and also between the repeat and TSR region are also indicated (5,6 and 7).

**Figure 3.**
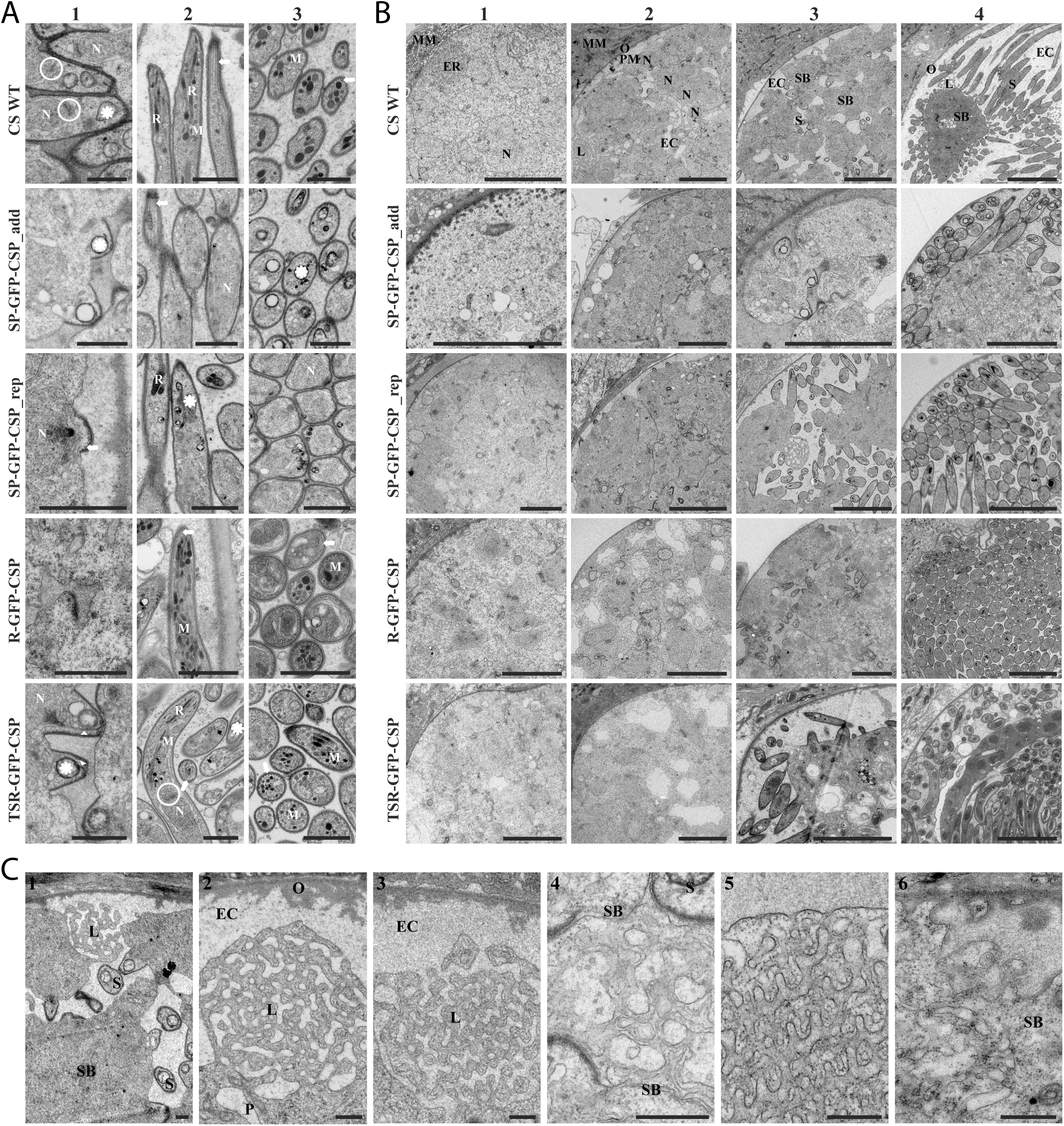
Electron micrographs show normal sporozoite development. Detail (A) and overview (B) transmission electron micrographs from oocysts at different stages of development from the different parasite lines. A: Sporozoites during the process of budding and in longitudinal and cross sections of the indicated parasite lines. Note the rhoptry Anlagen (asterisks), nuclei (N), micronemes (M), rhoptries (R) and microtubules (arrow) as well as the thickened pellicle (caused by the underlying IMC) during early budding. Arrowheads indicate possible rootlet fibers linking the apical tip and the nucleus. ER exit sites are always located at the apical end of the nucleus (ring). Scale bars: 1 µm. B: Oocysts from the different indicated parasite lines shown as quarters from oocyst wall to oocyst center to optimize overview while preserving detail. Early to late oocysts are shown from 1 to 4. MM: mosquito midgut, ER: endoplasmic reticulum, N. nucleus, O: oocyst wall, L: labyrinthine structure, EC: extracellular cavity, S: sporozoite, SB: sporoblast. Scale bars: 5 µm C: Labyrinthine structures (L) seen in wild type (1-3) and all generated parasite lines (4: SP-GFP-CSP_rep, 5: R-GFP-CSP, 6: TSR-GFP-CSP. EC: extracellular cavity, O: oocyst wall, S: sporozoite, P: plasma membrane, SB: sporoblast. Scale bar: 500 nm.

**Figure 4.**
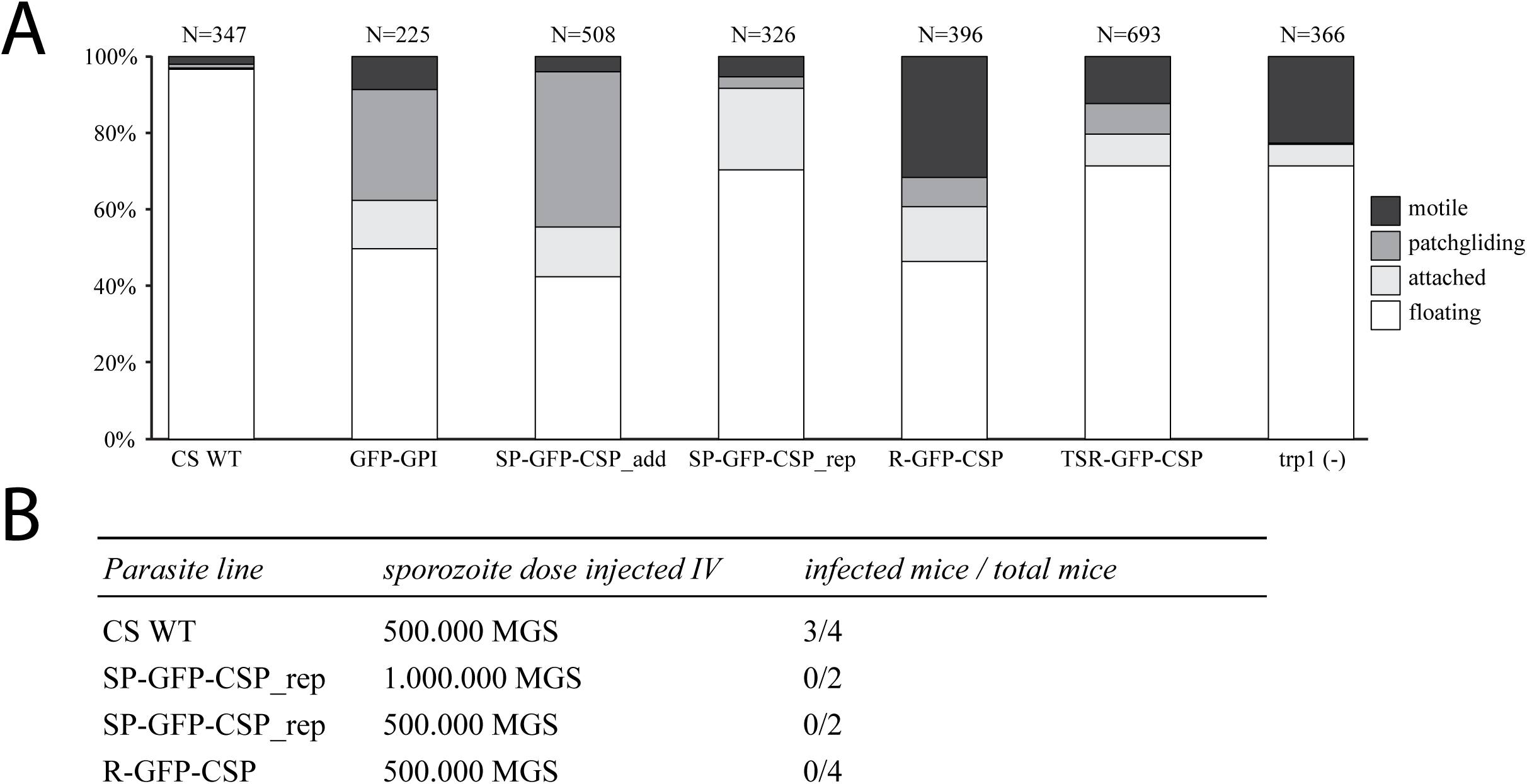
Gliding motility and infectivity to mice. A: Gliding motility of midgut derived CSP mutants. Midgut derived sporozoites were analyzed for gliding motility and classified into motile (> 30 μm traveled within 180 seconds), patch gliding (back and forward motion over a single attachment site), attached or floating. The number of analyzed sporozoites is indicate on top of the graph. Parasites lacking TRP1 (trp1 (-)) (Klug and Frischknecht, 2017) were analyzed for comparison. Samples were derived from mosquito midguts day 25 post infection and imaged after Accudenz purification. SP-GFP-CSP_rep was derived at day 18 post infection as they degenerated thereafter within the cysts. B: Infectivity of midgut sporozoites (day 22/23 post infection) inoculated intravenously into mice. Parasitemia was monitored from day 3 until day 20.

**Figure 5.**
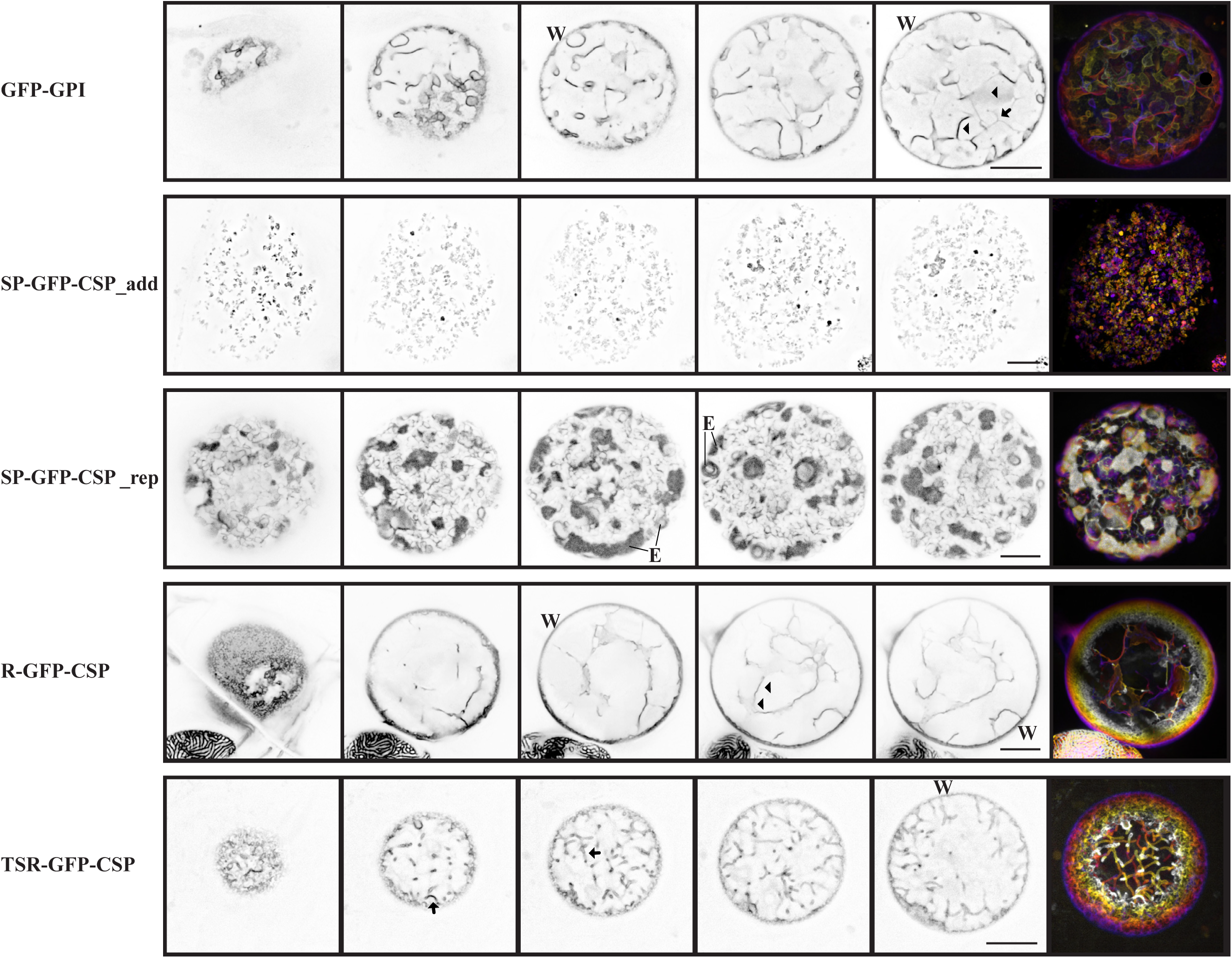
Early oocyst development. Tubular (arrows) and sheet-like (arrowheads) invaginations revealed during early invagination from the different indicated parasite lines. Orthogonally optically sectioned sheet-like invaginations show an apparent stronger signal. Individual panels show inverted fluorescent confocal sections and the merge shows a depth encoded 3D reconstruction of the z-stack. Oocyst of SP-GFP-CSP_add do not express GFP at the early oocyst stage, so Hoechst stained DNA is shown. Note the signal at (underneath) the oocyst wall (marked with W) and fluorescent accumulation in ER like structures (marked with E). Scale bars: 10 µm.

**Figure 6.**
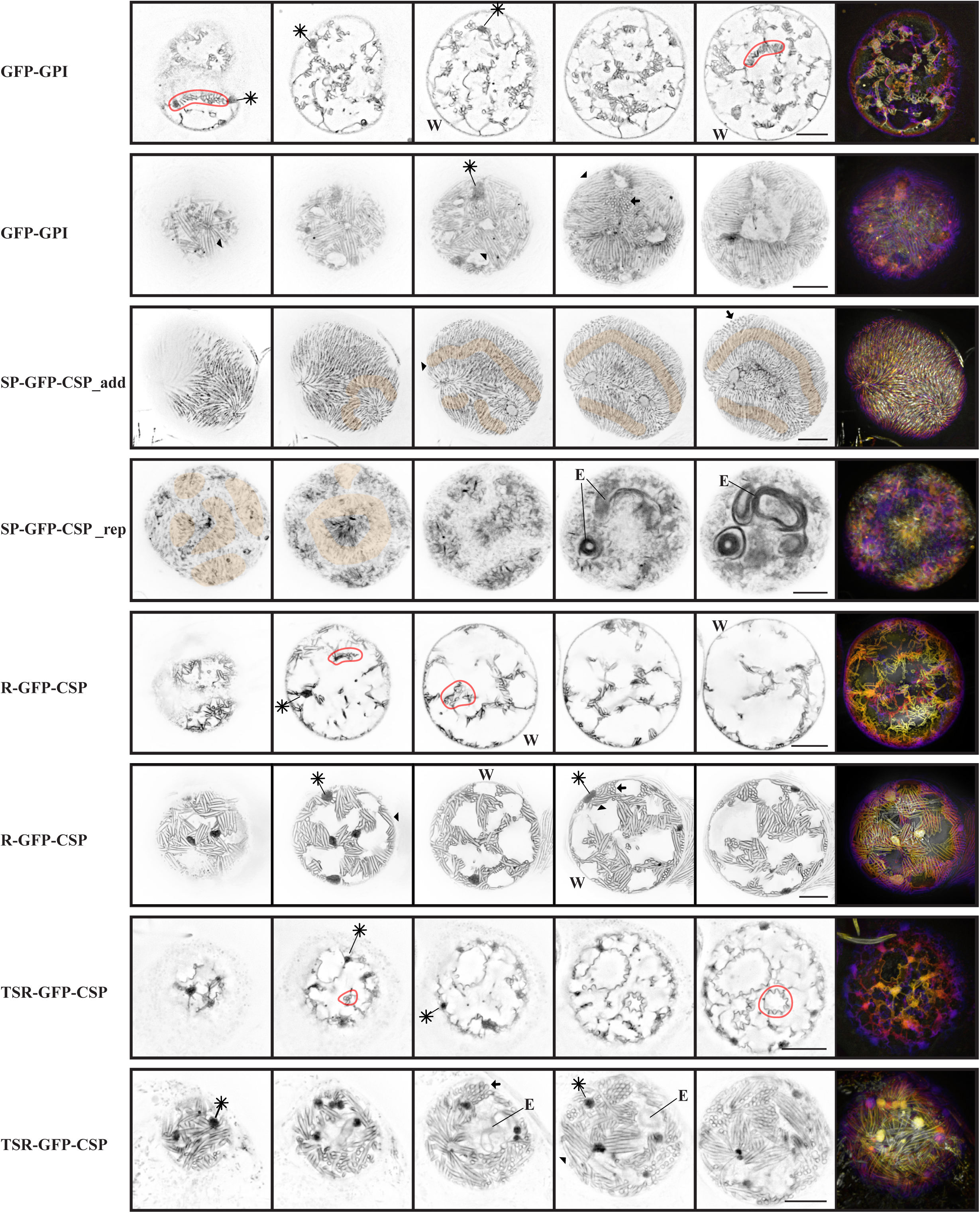
Oocysts during sporozoite formation. Early and advanced sporozoite formation from invaginations of the different indicated parasite lines. Individual panels show inverted fluorescent confocal sections and the merge shows a depth encoded 3D reconstruction of the z-stack. Note the cross (arrows) and longitudinal (arrowheads) sections through sporozoites and the non-membranous pattern in the SP-GFP-CSP parasite lines (nuclei of forming sporozoite indicated by orange shading). Large black dots (stars + line) likely correspond to labyrinthine structures. Note the signal at (underneath) the oocyst wall (marked with W) and fluorescent accumulation in ER like structures (marked with E). Exemplary areas of initiation of sporozoite apical tip formation resulting in plasma membrane curvature are circled in red. Scale bars: 10 µm.

**Figure 7.**
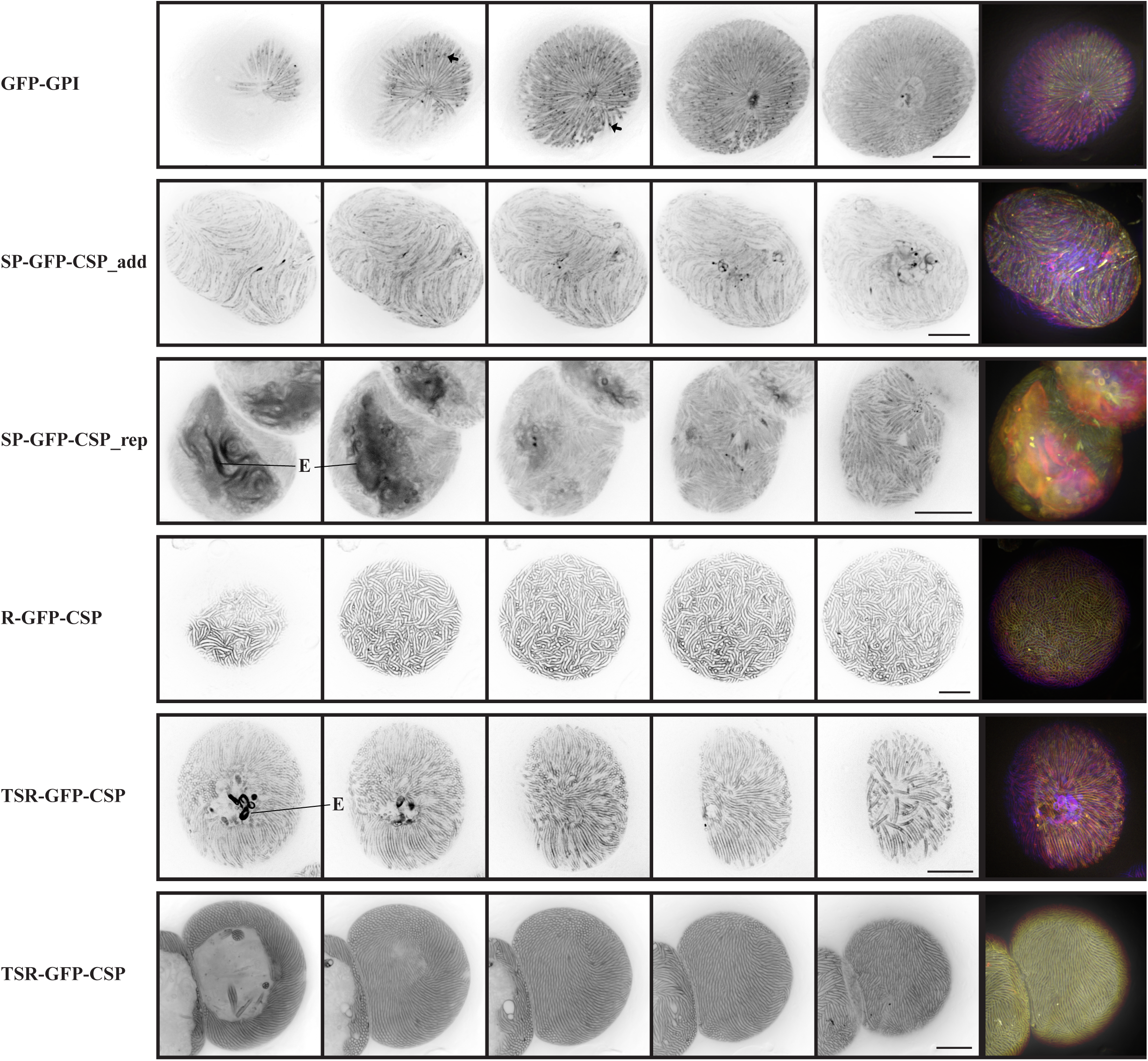
Oocysts with mature sporozoites. Fully formed sporozoites nearly fill the entire oocyst. Individual panels show inverted fluorescent confocal sections and the merge shows a depth encoded 3D reconstruction of the z-stack. Arrows point to examples of accumulations of signal within the GFP-GPI sporozoites. Note the different pattern in the SP-GFP-CSP parasite lines and that the SP-GPF-CSP_rep example shows a not completely developed oocyst with fluorescent accumulation in ER like structures (marked with E). Scale bars: 10 µm.

### Processing of GFP-CSP fusion proteins

To investigate if the fused proteins were indeed GFP-CSP fusions we examined western blots with antibodies against GFP and the repeats of CSP (Figure 2A). The anti-repeat antibodies recognized the known CSP double bands in GFP-GPI and SP-GFP-CSP parasite lines corresponding to the full-length protein and one lacking the N-terminus (Coppi et al., 2011). In contrast, the same antibodies recognized two additional higher bands in the R-GFP-CSP and TSR-GFP-CSP parasite lines suggesting that the latter two contain full length GFP-CSP fusion proteins as well as the N-terminally cleaved CSP-GFP version (Figure 2A and B). The anti-GFP antibody revealed two bands in parasites expressing GFP-GPI, indicative of the presence of both soluble GFP (low band) and GPI attached (higher band) forms or two forms of GFP-GPI that migrate at slightly different height. In the SP-GFP-CSP line no full length CSP-GFP could be detected with either anti-GFP or anti-repeat antibodies. A large GFP band was observed with a minor band just on top suggesting that GFP is cleaved very early after protein synthesis. In the R-GFP-CSP parasite line a major 37 kDa band could be detected with the anti GFP antibody suggesting the presence of a fusion protein including the GFP and the TSR. Also, a faint band was detected at 70 kDa, likely corresponding to GFP-CSP protein lacking the N-terminus as revealed by the anti CSP antibody (Figure 2A). For the TSR-GFP-CSP parasites anti-GFP antibodies identified multiple bands suggesting that some full length GFP-CSP protein is present but that proteolytic processing leads to multiple GFP-CSP species in these parasites (Figure 2B).

### Formation of sporozoites proceeds like in wild type parasites

We next investigated sporozoite formation by electron microscopy. This showed that all parasite lines developed normally in a manner reminiscent of wild type parasites (Figure 3A, B). Bud formation started after the nuclei aligned near the plasma membrane and coincided with the appearance of the thick pellicle due to the formation of the inner membrane complex during budding (Terzakis, Sprinz & Ward, 1966; Terzakis, Sprinz & Ward, 1967; Ferguson et al., 2014). Early sporozoite buds showed the typical round large vesicular rhoptry Anlagen. Sporozoites elongated thereafter and showed the typical elongated rhoptries, micronemes and microtubules. Finally, in all stages fully formed sporozoites could be detected in late stage oocysts (Figure 3A, B). We also noticed the labyrinthine structures, which represent a highly organized membranous structure that increase the surface area of the plasma membrane. Some of these also contain visible internal membranes (Figure 3C) (Wong and Desser, 1976; Meis, Verhave, Jap & Neuwissen, 1985).

### Motility of midgut sporozoites

Midgut sporozoites show only a weak capacity for surface adhesion which is a necessity for gliding motility (Vanderberg, 1974; Hegge et al., 2010; Hegge et al., 2012). Hence few midgut sporozoites are usually observed actively migrating on a substrate. Yet, sporozoites residing for longer than wild type parasites within the oocysts increase their capacity to glide (Klug and Frischknecht, 2017; Aly and Matuschewski, 2005) possibly reflecting a continued maturation process (Sato, Montagna & Matuschewski, 2014; Silva et al., 2016). We found that sporozoites from most lines were attaching more robustly at day 25 post infection than wild type sporozoites (Figure 4A). Typical back-and-forth movement termed patch-gliding (Münter et al., 2009) of hemolymph sporozoites was most frequently observed in GFP-GPI and SP-GFP-CSP_add parasites, while R-GFP-CSP sporozoites were gliding in the most robust manner. With over 20% gliding midgut sporozoites this reached the levels of hemolymph sporozoites (Münter et al, 2009; Klug et al., 2020). Over 10% of TSR-GFP-CSP sporozoites were also gliding. SP-GFP-CSP_rep parasites showed a motility pattern most similar to wild type, but with increased adhesion. These data suggested to us that R-GFP-CSP and SP-GFP-CSP_rep midgut sporozoites might be infective for liver cells. Strikingly, however, when we injected 500.000 R-GFP-CSP and 500.000 or one million SP-GFP-CSP_rep midgut sporozoites intravenously into mice, none developed a blood stage infection, while three out of four mice become blood stage patent after injection of 500.000 wild type midgut sporozoites (Figure 4B). This suggests, that like in the region II+ mutation (Wang et al, 2005), our GFP-CSP lines are not capable to enter liver cells.

### R-GFP-CSP and TSR-GFP-CSP but not SP-GFP-CSP localize to the plasma membrane

Investigation of the fluorescence signal during sporozoite formation using spinning disc confocal microscopy of acutely dissected mosquito guts showed that the fluorescent signal of all parasite lines with the exception of the two SP-GFP-CSP lines could be found at the plasma membrane of the oocyst, delineating the invagination of the plasma membrane, a process that precedes sporozoite formation (Figure 5). In the SP-GFP-CSP lines the signal was not found at the plasma membrane but was also distinct from cytoplasmic GFP. It likely stays within a membrane-delimited compartment and in early oocysts looks very similar to the endoplasmic reticulum accumulation observed in liver stages (Kaiser et al., 2016) (Supplementary movie S1). In late oocysts, whirl like accumulations of unclear origin are observed within the sporoblast center, which could correspond to a specialized organization of the endoplasmic reticulum (Figure 6, Supplementary movie S2). Later in sporozoite formation, the induction of plasma membrane curvature of budding sporozoites was readily observable in the R-GFP-CSP, TSR-GFP-CSP and GFP-GPI expressing parasites (Figure 6, Supplementary movie S3). Sporozoite budding could be followed until sporozoite formation was complete. In TSR-GFP-CSP the plasma membrane was strongly stained but weak GFP fluorescence was also observed from the ER, potentially from slowed or delayed trafficking (Figure 6, Supplementary movie S4).

In all lines but the two SP-GFP-CSP ones strong fluorescence accumulations could be detected in literally all imaged cysts, albeit not in every optical section (Figure 6, Supplementary movies S3-5). In contrast to the ER accumulations observed in SP-GFP-CSP_rep that were observed earlier, these signals were only observed with the onset of apical tip formation, mostly bridging two sporoblasts. They likely represent the highly convoluted membranous structures previously named labyrinthine structures (Meis, Verhave, Jap & Neuwissen, 1985). Their apparent diameter in the fluorescent images was around 1-5 µm (n=53), while those in the EM images measured from 0.5-4 µm (n=13). In two dimensions, these structures are sponge like in appearance by EM (Figure 3). Roughly 50-70% of the volume is cytoplasmic content and internal membranous structures, the rest extracellular volume. The distance between opposing plasma membranes is between 200-400 nm, and in some instances the substructures represent a short comb. Finally, in all parasite lines fully developed sporozoites could be detected. Sporozoites in fully matured oocysts labelled similar to those found in isolated sporozoites (Figure 7 – compare with Figure 1C, Supplementary Movie S6). In late oocysts of the SP-GFP-CSP_rep line a number of unusual structures were found with the GFP signal appearing localized to a membranous structure. In these oocysts few normal sporozoites could be detected, suggesting that less cytoplasmic material could be converted into sporozoites (Figure 7, Supplementary Movie S2). This in turn might also explain why midgut sporozoites of this line were not infectious if injected into mice (Figure 4B).

### Microtubules appear after initial bud formation during early sporogony

We recently succeeded in labeling microtubules with SiR-tubulin within oocysts (Spreng et al., 2019). SiR tubulin intercalates with microtubules and only then becomes fluorescent, hence allowing selective staining of microtubules in live cells (Lucinavicius et al., 2014). Using this dye in combination with our parasite lines and Hoechst for labeling of nuclei showed that microtubules were absent from oocysts with smooth invaginated plasma membrane even if nuclei were already recruited to the periphery (Figure 8A, B, first row). Only in oocysts with signs of early apical bud formation could we detect microtubules, suggesting that they only form after the initiation of sporozoite budding (Figure 8).

**Figure 8.**
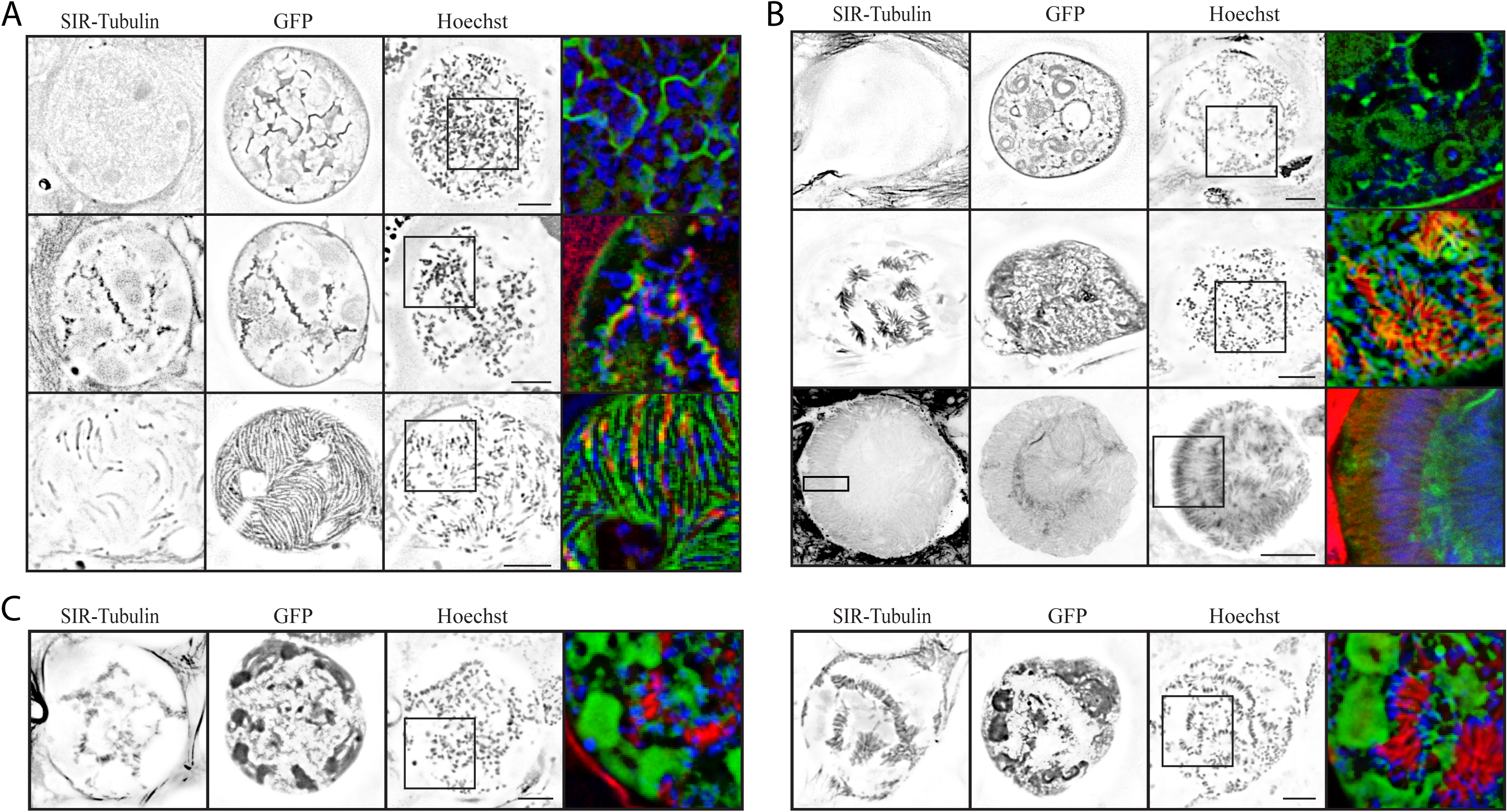
Microtubules form after initial bud formation. A-C: Example images of confocal sections at different stages of selected parasites lines show microtubules (red – SiR tubulin), nuclei (blue – Draq594) and the indicated GFP-CSP fusions. Note that nuclei align at the plasma membrane prior to microtubule assembly, while microtubules are found at highly curved membranes corresponding to budding sporozoites. Scale bar: 10 µm A: Oocysts of R-GFP-CSP. B: Oocysts of TSR-GFP-CSP. Last row: fixed and stained for tubulin with anti-Tubulin antibody. C: Oocysts of SP-GFP-CSP_rep parasite line.

**Figure 9.**
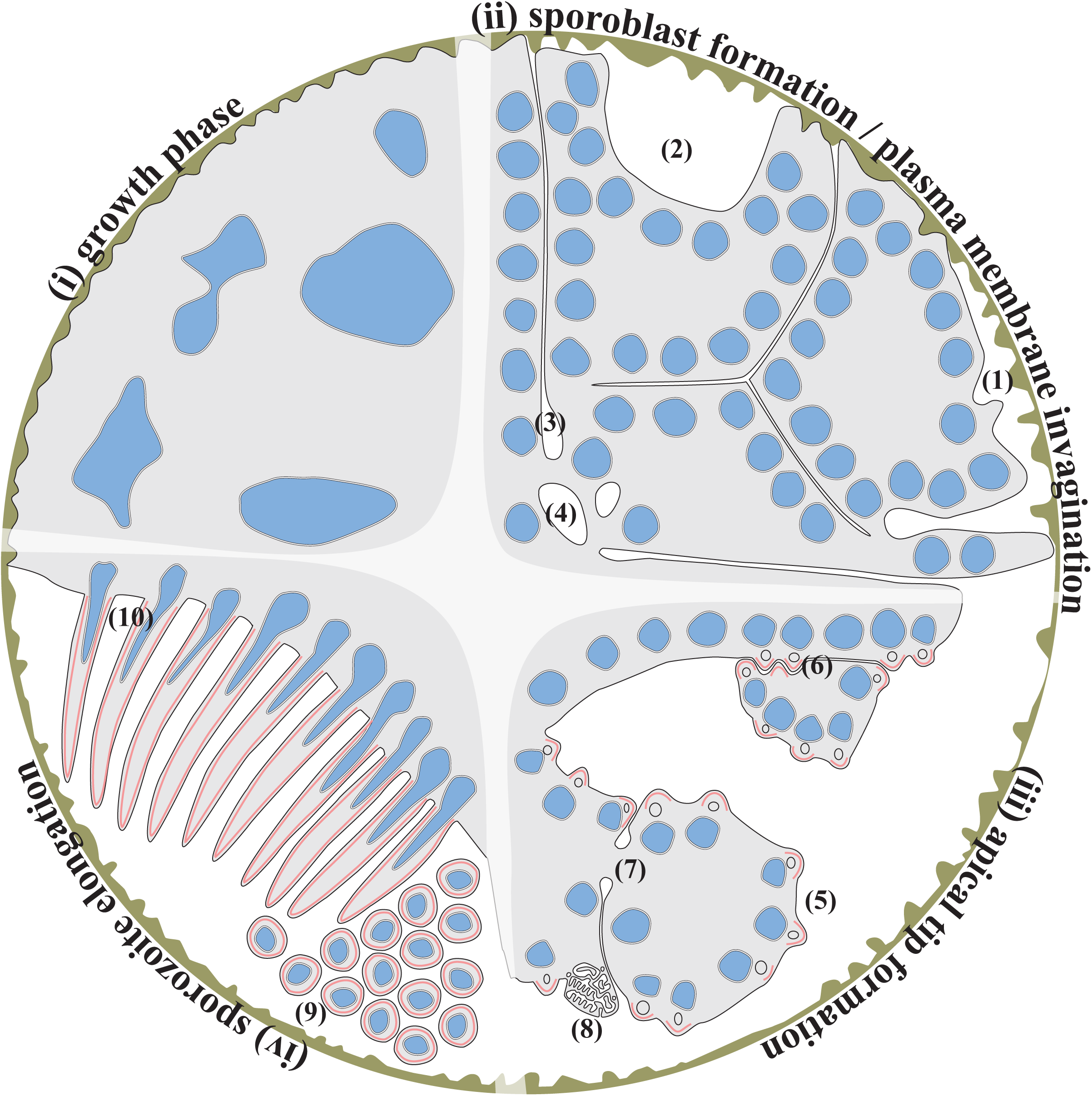
Model of Oocyst development. Oocyst development can be separated into four phases. (i) Growth phase: After rounding up of the ookinete a strong increase in cell size, genome replication and oocyst wall (brown) formation occurs. (ii) Plasma membrane invagination and sporoblast formation: nuclear division to many and small nuclei (blue) coincides with the retraction of the plasma membrane (PM) (black) from the oocyst wall (1 and 2) and deep invaginations of the PM (3). The nuclei collect underneath the PM. Internal membrane structures spread towards and fuse with the PM (4). (iii) Apical tip formation: Between the PM and the underlying nuclei the apical tip of forming sporozoites is initiated, visible by formation of inner membrane complex (IMC) directly accompanied by microtubules (together shown in red). This is followed by the appearance of the rhoptry Anlagen (circle) and a bulging PM. This can occur outside of sporoblasts (5) as well as in between sporoblasts (6). Note the cytoplasmic bridges in between sporoblasts (7) and labyrinthine structures at the periphery of sporoblasts, frequently located in between them (8). (iv) Sporozoite elongation: During the last stage of sporozoite development prior to egress, sporozoites elongate by uniform retraction of or pushing through the PM. The “end” of the forming IMC and microtubules (red) always coincides with the deepest PM invagination. At this time sporozoites are in similar orientation than sporozoites in their local environment (9 and 10) until they separate from the remaining sporoblast.

## Discussion

### Oocyst egress is blocked in all fusions but SP-GFP-CSP and GFP-GPI

We generated a series of *P. berghei* parasite lines expressing different fusions of the green fluorescent and circumsporozoite proteins. We designed these as internal GFP fusions with the GFP placed between known domains of CSP. We hoped that one of these would allow normal progression of sporozoites along their long journey from formation in the oocyst to de-differentiation in hepatocytes. While the generated lines allowed observation of sporozoite formation, they either cleaved off the GFP if placed right after the signal peptide, or arrested after maturation within oocysts. The latter, literally dominant negative impact of the fusion protein might indicate that the function of CSP in oocyst egress (Wang et al., 2005) is compromised by the presence of the bulky GFP. Although it is not clear how CSP mediates sporozoite egress it is possible that it needs to be proteolytically cleaved and that hence processing is modulated in the GFP-CSP fusions. This could also explain why midgut-derived CSP-GFP sporozoites are not infective for rodent livers, where processing of CSP was shown to be induced (Coppi et al., 2011). Alternatively, removal of the GPI-anchor by phospholipases might also be blocked by the GFP (Hereld et al., 1986). Indeed, our western blots detect many different bands which cannot easily be accounted for but hint towards complex functions of the protein. But why do SP-GFP-CSP_rep mutants not get out of the oocysts? In these mutants all CSP is generated as GFP-fusion, but the GFP is cleaved before CSP arrives at the plasma membrane. Yet these sporozoites do not egress. Post signal peptide N-terminal processing of CSP might be due to the two N-terminal PEXEL motifs found in CSP (Figure S1) (Singh et al., 2007). However it would be expected that a PEXEL motif after GFP would not be recognized, at least indicated by PEXEL motif processing in *Plasmodium falciparum* bloodstages (Hiss et al., 2008). This would result in a modified N-terminus of SP-GFP_CSP_rep compared with CSP WT. Clearly more work is needed to understand the possibly many different cleavage events necessary for CSP function.

### Trafficking of CSP

The observation by fluorescence microscopy that GFP accumulates within internal structures in the SP-GFP-CSP sporozoites and the nearly complete cleavage of this fusion protein as determined by Western blot, suggests that GFP is cleaved off already within the ER or only shortly after. Intriguingly, in sporozoites showing CSP-GFP fusions at the PM, the protein is almost exclusively found at the PM. This is in contrast to GFP-TRAP, which is stored in micronemes (Kehrer et al., 2016). These observations hint that CSP is trafficked differently prior to GFP maturation, i.e. not within micronemes and faster than TRAP. Interestingly, the short fusion protein of GFP and the GPI-anchor of CSP shows some internal labeling (Figure 1C) that is different in localization to the apically located GFP-TRAP (Kehrer et al., 2016) and within the proximity of the ER exit site and Golgi. In direct comparison on trafficking, R-GFP-CSP is always detected only at the PM, whereas TSR-GFP-CSP showed a faint fluorescence within the ER of oocysts (Figure 6). This indicates that the presence of GFP directly adjacent to the GPI-anchor might slightly interfere with trafficking, especially if the GPI-anchor itself is the signal for trafficking as is the case in *Trypanosoma brucei* (Kruzel, Zimmett 3rd & Bangs, 2017; Triggs and Bangs, 2003). Comparing the relative intensity of the full length bands with the multiple smaller bands detected with anti-GFP antibody suggests that a substantial fraction of R-GFP-CSP as well as TSR-GFP-CSP protein is not full length. Our data does not allow to conclude whether the protein is processed en route or only after it appears on the surface, but the fact that no internal structures are labelled in this parasite line would favor processing upon arrival at the PM.

### Labyrinthine structures

In almost every section of every oocyst observed with EM we found labyrinthine structures, which are small prior to the start of sporozoite budding and most prominent during sporozoite budding and disappear/disintegrate once sporozoite elongation is completed. Although we did not perform correlative microscopy here, several observations suggest that the bright fluorescent dots in GFP-GPI, R-GFP-CSP and TSR-GFP-CSP correspond to labyrinthine structures. First, their size is similar in both EM and fluorescence microscopy. Secondly, we could not observe any fluorescence dots in the SP-GFP-CSP parasite lines. Thirdly, their time of appearance and localization at the proximity of sporoblasts match between EM and fluorescence microscopy. But what are their function?

The strong fluorescent intensity of the labyrinthine structures in comparison to the PM of oocysts suggests that the main membrane component is PM that accumulates at high density. Indeed, EM shows a highly convoluted layer of membranes (Figure 3C). EM also shows an internal membrane network. The origin of this is completely unknown. Structures with very similar appearance have been observed previously before merozoite formation in liver schizonts, and proposed to be involved in nutrient uptake (Meis et al., 1985). Structures which are similar to the rather degenerate labyrinthine structures observed in old oocysts of TSR-GFP-CSP oocysts have been described in oocysts of *Leucocytozoon dubreuli* as ER-associated vesicles (Wong et al, 1976). In the parasites lacking the CSP repeats degraded labyrinthine structures could also be observed, but these were not commented on by the authors (Ferguson et al, 2014). However, we did not observe any labyrinthine structures prior to sporoblast development by PM invagination. This makes the function as a general nutrient uptake area less likely as nutrients are likely most needed during the growth of oocysts. Yet, nutrient uptake might be blocked by the formation of the IMC and hence these structures are established before the entire PM is covered with IMC during sporozoite formation. Additionally or alternatively, this structure could be involved in the secretion of CSP itself. However CSP is secreted prior to appearance of the labyrinthine structures. In trypanosomes, endocytic recycling of the GPI-anchored protein VSG has been described in detail (Engstler et al., 2004). Potentially these structures are involved in protein sorting by size, where the short membrane form is replaced by the full length CSP detected in salivary gland sporozoites. If protein size is the main sorting mechanism, this could also give another explanation why this fails in GFP-CSP fusions. Another study has recently been performed on a PM localized transmembrane protein that has been tagged with GFP (Burda et al., 2017) but it is not clear if this also localizes to the labyrinthine structures.

### Towards 4D imaging of oocysts in mosquitoes

Recent advances in light microscopy enable imaging of complex and large structures in 3D over time (4D). The use of confocal microscopy as shown here could in principle be extended to live imaging by repeatedly imaging oocysts. Combining plasma membrane labels with nuclear or organelle-specific labelling could hence allow the visualization of sporozoite formation and answer questions such as whether organelle packaging is highly spatially or temporally coordinated or even whether the organelles are somehow coupled to each other. Furthermore, the use of truncated proteins fused with GFP could yield insights into where these proteins function during sporogony. Two studies along these lines have shown that variations of CSP or tubulin protein expression levels can hamper sporozoite formation and impact their form, motility and infectivity (Thathy et al., 2002; Spreng et al., 2019). Here we used SiR tubulin to show that microtubules only form once sporozoite bud formation has proceeded in fixed images (Figure 8). Using 4D microscopy should allow to determine the precise timing.

In conclusion, we generated a series of parasite lines expressing different fusion proteins of GFP and CSP. This showed that only internal GFP-tagging allowed the detection of GFP-CSP fusion proteins at the plasma membrane. The introduced additional copies of correctly localized GFP-CSP fusion proteins stopped egress from oocysts. Hence to enable imaging of CSP in sporozoites during transmission from mosquito to mammal these GFP-CSP fusion proteins should be expressed from a promoter that is active only after sporozoites entered salivary glands. The observation that some GFP-CSP fusion proteins were not full-length suggest complex processing to occur, some of which is likely interfered with by the generated fusions.

## Methods

### Animal work

All mice experiments were performed according to the FELASA and GV-SOLAS standard guidelines and were approved by the German authorities (Regierungspräsidium Karlsruhe). Parasite generation and maintenance was performed in NMRI mice and sporozoite injections were performed with female C57BL/6 mice (both from Charles River).

### Bioinformatics

All genetic sequences were retrieved from PlasmoDB (https://plasmodb.org/plasmo/, release 6.4-30) and GeneDB (www.genedb.org/Homepage) (Aurrecoechea et al, 2008; Logan-Kumpler et al, 2012). Sequence alignments were performed with clustalW2 (https://www.ebi.ac.uk/Tools/msa/clustalw2/) and CLC Workbench 7.9.1 (CLC bio, Qiagen bioinformatics, USA) and manually curated. Signal peptide prediction was performed with Signal IP 4.1 (http://www.cbs.dtu.dk/services/SignalP/), GPI-anchor prediction was performed with PredGPI (http://gpcr.biocomp.unibo.it/predgpi/) (Petersen, Brunak, von Heijne & Nielsen, 2011; Pierleoni, Marelli & Casadio, 2008). Plasmid design was performed with ApE v2.0.45 (https://www.thejorgensenlab.org/).

### Generation of parasite lines

Mapping of the exact position for internal GFP tagging was performed upon sequence alignment and predicting the signal peptide cleavage site and GPI-anchor prediction (Figure S1). Primers were obtained from Thermo Fischer Scientific (For sequences see Figure S3), restriction enzymes from New England Biolabs. All required *Plasmodium* sequences were amplified from genomic DNA from *Plasmodium berghei* strain ANKA with high fidelity Taq polymerase (Thermo Fischer Scientific, Waltham, USA) with 8 °C lowered elongation temperature and verified by Sanger sequencing (GATC; now Eurofins, Konstanz) upon cloning.

For generation of GFP-GPI, the promoter region including the SP was amplified with primers P208 and P268, and cloned via EcoRI and NdeI into Pb238 (Singer et al., 2015). Then the GPI-anchor sequence as well as a short 3’UTR was amplified with P274 and P270 and cloned via KasI and EcoRV. The resulting vector was digested with PvuI and integrated via double crossover into a silent chromosome 12 locus (Figure 1B).

For interdomain tags, the promoter region up to the respective *gfp* insertion site was amplified with P208;P268 for SP-GFP-CSP, P208;P271 for R-GFP-CSP and P208;P273 for TSR-GFP-CSP and cloned via EcoRI and NdeI into Pb238. The remaining piece of *csp* with the 3’UTR was amplified with P269;P270 for SP-GFP-CSP, P272;P270 for R-GFP-CSP and P274;P270 for TSR-GFP-CSP and cloned via KasI and EcoRV. Constructs were linearized with PacI (SP-GFP-CSP), PmlI (R-GFP-CSP and TSR-GFP-CSP) and integrated via single crossover into the *csp* locus (Figure 1B).

Transfection was performed in *P. berghei* strain ANKA as published (Janse, Ramesar & Waters, 2006). Transgenic parasites were selected with pyrimethamine via the drinking water (0.07 mg/ml) in NMRI mice. Upon successful initial genotyping, parasites were cloned by limiting dilution of 0.7 parasites injected *i*.*v*. into 10 NMRI mice. All generated clones were genotyped again.

For genotyping, 5’ integration and 3’ integration was probed with P134;P210 P137;P99 (GFP-GPI), P267;P210 P893;P882 (SP-GFP-CSP, R-GFP-CSP, TSR-GFP-CSP) and P267;P210 P234;P882 (SP-GFP-CSP_rep). The whole locus was amplified with P134;P137 (GFP-GPI) and P267;P882 (all others). Expected sizes are indicated in Figure S2. Note that the whole locus was not always obtained due to its large size.

### Mosquito work

*Anopheles stephensi* mosquitos (strain SDA 500) were infected as described previously (Frischknecht et al, 2004). Parasite development was monitored from day 10 post infection by dissection. Mosquitoes were washed in 70% ethanol and stored in PBS and dissected under a Binocular Nikon SMZ 1500 with GFP illumination for preparation of midgut and salivary gland samples. For haemolymph preparation sporozoites were dissected ‘dry’: The last two segments of the abdomen were removed, the tip of a self-made glass capillary was inserted into the spiracle of the mesothorax and the haemolymph was rinsed with PBS. Subsequently the midgut and salivary glands were dissected from the same mosquito.

### Light microscopy

Imaging was performed on an inverted Axiovert 200 M microscope from Zeiss, a spinning disc confocal ERS-FRET from PerkinElmer using a Nikon inverted microscope or Leica SP5 confocal microscope. Gliding assays were performed in RPMI with 3% BSA. Sporozoites from midgut samples were purified using 17% Accudenz gradient centrifugation (Kennedy et al., 2012). For IFA, midguts were fixed in 4% PFA for 30 minutes, permeabilized ON with 0,5% Triton-X-100 with 3% BSA, incubated overnight with anti-Tubulin antibody. Secondary antibody Alexa Fluor 546 (Invitrogen, Karlsruhe, Germany) was incubated overnight together with Draq5 (Thermo Fischer Scientific, Waltham, USA) to label DNA and washed 5 times for 20 minutes and mounted with Prolong Gold (Thermo Fischer Scientific, Waltham, USA).

Life cell microscopy of whole midguts was performed in RPMI with 3% BSA. For tubulin staining, midguts were incubated for 30 minutes with 1 μM SIR-Tubulin (Spirochrome, Stein am Rhein, Switzerland) (Lukinaviecius et al., 2013) as well as 1 μg/ml of Hoechst 33342 (Thermo Fischer Scientific, Waltham, USA), washed in fresh RPMI with 3% BSA and sealed with a 1:2:1 mixture of lanolin:paraffin:vaseline.

### Image processing

Image analysis was performed with FIJI (LOCI, Wisconsin-Madison, USA) (Schindelin et al., 2012). Figures were generated with Illustrator CS5.1 software (Adobe, München, Germany) and Photoshop CS 5.1 software (Adobe, München, Germany). Images shown in Figure 8 were deconvolved with Autoquant X3 software (Media Cybernetics). Multiple optical sections shown in Figure 5, 6 and 7 where contrast adjusted for each individual slice to improve structural information. If required, the intensity spectrum of the image was collapsed into 8 bit by square rooting the entire image.

### Western blotting

Dissected sporozoite samples were purified with 17% (w/v) Accudenz gradient centrifugation. Samples were lysed with freshly prepared ice-cold RIPA buffer with protease inhibitor (Roche, Mannheim, Germany) for one hour on ice. Samples were separated in 4-15% precast gels and transferred semi-dry using the BioRad Transblot turbo system. Samples were incubated with primary antibodies; anti-GFP antibody 13.1 + 7.1 (Roche, Merck, Darmstadt, Germany) or anti-CSP repeat antibody – mAB 3D11 (Yoshida et al., 1980) and HRP bound antibodies (GE healthcare, Thermo Fischer Scientific, Waltham, USA) and incubated with SuperSignal West Pico chemiluminescent solution (Thermo Fischer Scientific, Waltham, USA).

### Electron microscopy

Electron microscopy was performed at the Electron Microscopy Core Facility (EMCF) of Heidelberg University. Midguts were dissected directly into the fixation buffer and sample preparation was performed by the core facility technician Steffi Gold, using classical chemical fixation. Primary fixation was performed in 2% glutaraldehyde with 2% PFA in 100 mM sodium cacodylate buffer at 4 °C overnight. Sample was washed 3 times with 100 mM caco buffer and secondary fixation was performed with 1% osmium in 100 mM caco buffer for 60 min at room temperature. Sample was washed twice in caco buffer, twice in dd H_2_O and contrasted in 1% uranylacetate in dd H_2_O at 4 °C overnight. Samples were washed in dd H_2_O twice and dehydrated in 30%, 50%, 70%, 90%, 100% and 100% acetone for 10 minutes each. Sample was then embedded in Spurr resin (23,6% epoxycyclohexylmethyl-3,4-epoxycyclohexylcarboxylate (ERL), 14,2% ERL-4206 plasticizer, 61,3 nonenyl succinic anhydride, 0,9% dimethylethanolamine) and for this incubated in 25%, 50% and 75% 45 min at room temperature each and in 100% at 4 °C overnight. Embedding was finalized in BEEM capsules overnight at 60 °C. Samples were trimmed and cut into 70 nm thick sections. Electron microscopy was performed on a Joel JEM-1400 transmission microscope with a bottom mount 4k digital camera (F416, Tietz Video and Image Processing Systems GmbH, Gauting) with the assistance of Dr. Stefan Hillmer.

## Acknowledgements

We thank Rogerio Amino, Amanda Balaban, Carolina Thieleke-Matos and Photini Sinnis for fruitful discussions and comments on the manuscript, Markus Meissner for support, Miriam Reinig for mosquito production, Catherine Moreau for help with cloning and Jannik Traut for help with microscopy. The work was funded by the Human Frontier Science Program (RGY0071/2011), the Deutsche Forschungsgemeinschaft (DFG, German Research Foundation) – project number 240245660 – SFB 1129 and the European Research Council (ERC StG 281719). FF is a member of CellNetworks cluster of excellence at Heidelberg University. We acknowledge the microscopy support from the Infectious Diseases Imaging Platform (IDIP) at the Center for Integrative Infectious Disease Research and thank Stefan Hillmer and Stephanie Gold from the Electron Microscopy Core Facility of Heidelberg University for support and the use of their microscopes.

