## Supplemental Figures for "Fluorescent tagging of *Plasmodium* circumsporozoite protein allows imaging of sporozoite formation but blocks egress from oocysts"

**Supplementary Figure 1** Alignment of circumsporozoite protein.

The amino acid sequence of CSP from different *Plasmodium* species was aligned with clustalW and manually curated to highlight conserved features of CSP. Conserved residues are colored according to type; green = hydrophobic, red = acidic, blue = basic, orange = polar, pink = prolin and glycine, yellow = cysteine. The two Pexel motives and the predicted N-terminus after signal peptide cleavage are highlighted in bold. Region I is emphasized with a box, basic residues implicated in hepatocytes invasion adjacent to Region I are in bold (Zhao, Bhanot, Hu & Wang, 2016). Perfect repeats are shown only once and copy number is indicated. The glycosylation motif WXXC is highlighted with a box (Swearingen et al., 2016). The amino acid that is GPI-anchored, the ω-site, is indicated in bold red, followed by the residues ω+1 and ω+2, highlighted in bold. The anchor addition signal also consists of the flexible and the hydrophobic region, which are cleaved upon GPI-anchor addition. GFP insertion sites are highlighted. GFP insertion for SP-GFP-CSP results in the signal peptide cleavage site shifted to CQG – GG (glycine linker of GFP). GFP insertion for TSR-GFP-CSP results in GPI-anchor addition to the second last C-terminal acid of GFP (DELYKG – GPI).

**Supplementary Figure 2** Genotyping of GFP-CSP strains.

Genotyping of gDNA was performed for all CSP mutants as well as ANKA WT gDNA. Expected sizes of PCR products are indicated below the gel images. GFP-GPI as well as SP-GFP-CSP_add parasites are not clonal and may contain low level of WT. For all others, genotyping after limiting dilution is shown. Some whole locus (WL) PCR products could not be amplified due to their large size. Unspecific PCR products are highlighted with a star.

**Supplementary Figure 3**

List of all Primers used in this study.

**Supplementary Movie S1**

Z-projection through an early oocyst of SP-GFP-CSP_rep. GFP signal is shown in green, DNA in blue.

**Supplementary Movie S2**

Z-projection through a late oocyst of SP-GFP-CSP_rep. GFP signal is shown in white.

**Supplementary Movie S3**

Z-projection through a early oocyst of SP-GFP-GPI. GFP signal is shown in green, DNA in blue.

**Supplementary Movie S4**

Z-projection through a oocyst of TSR-GFP-CSP. GFP signal is shown in white.

**Supplementary Movie S5**

Z-projection through a oocyst of R-GFP-CSP. GFP signal is shown in white.

**Supplementary Movie S6**

Z-projection through a late oocyst of SP-GFP-CSP_add. GFP signal is shown in green, DNA in blue.

```
signal peptide                 PEXEL      n-terminus
                                   [GFP]
Pb    CSP M----KKCTI LVVASLLLVN SLLPGYGQNK SIQAQRNLNE LCYNE-GNDN KLYHVL---- NSKNGKIYNR
Py    CSP M----KKCTI LVVASLLLVD SLLPGYGQNK SVQAQRNLNE LCYNE-ENDN KLYHVL---- NSKNGKIYNR
Pm    CSP M----KKLSV LAISSFLIVD FLFPGYHHNS NSTKSRNLSE LCYNNVD--T KLFNELEVRY STNQDHFYNY
Po    CSP M----RNLAI FAVSAFLFAD SRYPVYAHNG NSTEGRKLNE LCYNNVDLYN TLFNELDVGS STNQAFFLNS
Pg    CSP M----KKLAI LSASSFLFAD FLFQEYQHNG NYKNFRLLNE VCYNNMNI-- QLYNELEMEN YMSNTYFYNN
Prel  CSP MYFKMKKLAI LSTASFLFAD FLLQEYQHDA NYNKSRILNE VCYNNTNI-- QLYNELEKGS YESQTYFYNT
Pf    CSP M---MRKLAI LSVSSFLFVE ALFQEYQCYG SSSNTRVLNE LNYDNAGT-- NLYNELEMNY YGKQENWYSL
```

```
PEXEL                          n-terminus                                           RI
Pb    CSP NTVNRLLADA ---------- ---------- ---------- PEG------K KNEKK--NEK IERNNKLKQP
Py    CSP NIVNRLLGDA LNGKPEEKKD DPPKDGNKDD LPKEEKKDDL PKEEKKDDPP KDPKKDDPPK -EAQNKLNQP
Pm    CSP NKTIRLLNEN ---------- ---------- ---------N NEKDGNV--- TNERKKKPTK -AVENKLKQP
Po    CSP KKTLRLLNEN ---------- ---------- ---------- -PKE------ -NKKKRKDDK KPVENKLKQG
Pg    CSP KKTIRLLGEN -----DNEAN VNRANNNVAN DNRANGNRGN VNRA------ -NDRNIPYFR -ENVVNLNQP
Prel  CSP KKTHRLLGEN ---------- ---------- ---------- GNRVNEVIPI NNGRNNNGIK -NRRLNLRQL
Pf    CSP KKNSRSLGEN ---------- ---------- ---------- DDGNNE---- DNEK---LRK -PKHKKLKQP
```

```
                                  repeat region
Pb    CSP ------------ ------------ [PPPPNPND]4 - [PPPPNAND]2 - [PAPPNAND]4 - PPPPNPNDPAPP
Py    CSP VV--ADENVD-- ------------ [QGPGAP]21--- [QEPP]7 ----- ------------ ------------
Pm    CSP -----PGDDD-- --GAGNDAGNDA [GNAA]4 ------GNDA[GNAA]16- A[GNAA]3 GAA- -[GNAA]14GNE-
Po    CSP -----EREND-- ------------ [PPAPQGEGN]5 PPAAQGEGN--- [PPAAQGNGN]3 PPA---------
Pg    CSP V----------- ------------ [GGNGGVQPA]4 GGNGGAQPVAAG GGAQPVVADGGV QPLRQEGDAEED
Prel  CSP ----AE------ ------------ [GAGNGA]8 GAG ------------ ------------ ------------
Pf    CSP ----ADG----- ----NPDPNANP [NVDPNANP]3 - [NANP]16NVDP- [NANP]18NKNN- ------------
```

```
          repeat region                          linker
                                                    [GFP]
Pb    CSP NANDPPPPNPND PAPPQGNNNPQP QPRP[QP]9 --R PQPQPQPGG--- ------------ NNNNKNNNNDDS
Py    CSP ------------ ------------ -----------Q QPRPQPDG---- ---------NNN NNNNNGNNNEDS
Pm    CSP ------------ ------------ ------------ ----------KA KNKDNKVDANTN KKDNQEENNDSS
Po    CSP ------------ ------------ ------------ ----PAG----- -------KGKNE NQKEKEEKNAAN
Pg    CSP [GGNGGVQPA]11 [GGNGGAQPA]10 GGNDAAKPDGGN DDDKPEGGD--- ------------ EKSEEEKEDEPI
Prel  CSP ------------ ------------ ------------ ------------ ------------ ---DGKKEEKPI
Pf    CSP ------------ ------------ ------------ ----QGNGQGHN MPNDPNRNVDEN ANANSAVKNNNN
```

```
          RIII                  RII+                   remaining α-TSR
                                                                                 [GFP]
Pb    CSP YIPSAEKILE FVKQIRDSIT EE-WSQCNVT CGSGIRVRKR KGS-NKKAE- DLTL-EDIDT EICKMDKCSS
Py    CSP YVPSAEQILE FVKQISSQLT EE-WSQCSVT CGSGVRVRKR KNV-NKQPE- NLTL-EDIDT EICKMDKCSS
Pm    CSP NGPSEEHIKN YLESIRNSIT EE-WSPCSVT CGSGIRARRK VDAKNKKPA- ELVL-SDLET EICSLDKCSS
Po    CSP NPPSEDDIKK YIDKIRNDIT -ENWSPCSVT CGFGVRVRRK AGASAKKAQ- ELTL-SDLET EICKIENCSS
Pg    CSP PDPTQEEIDK YLKSILGNVT SE-WTNCNVT CGKGIQAKIK STSANKKRE- EITP-NDVEV KICELERCSF
Prel  CSP PDPTKEEIQK YINSVLDSVT SE-WSKCNVK CGKGVQVRIR PDS-NKNKGD KLTI-NDVEA KICELKRCSS
Pf    CSP EEPSDKHIKE YLNKIQNSLS TE-WSPCSVT CGNGIQVRIK PGSANKPKD- ELDYANDIEK KICKMEKCSS
```

```
          flexible   hydrophobic
Pb    CSP IFNIV-SNSL GFVILLVLVF FN
Py    CSP IFNIV-SNSL GFVILLVLVF FN
Pm    CSP IFNVV-SNSL GIVLVLVLIL FH
Po    CSP IFNVV-SNSL GLVIFLFLVF FH
Pg    CSP SIFNVISNSL GLAIILTFLF FY
Prel  CSP SIFNAISNSL GLAIFLVFLF FY
Pf    CSP VFNVVNS-SI GLIMVLSFLF LN
```

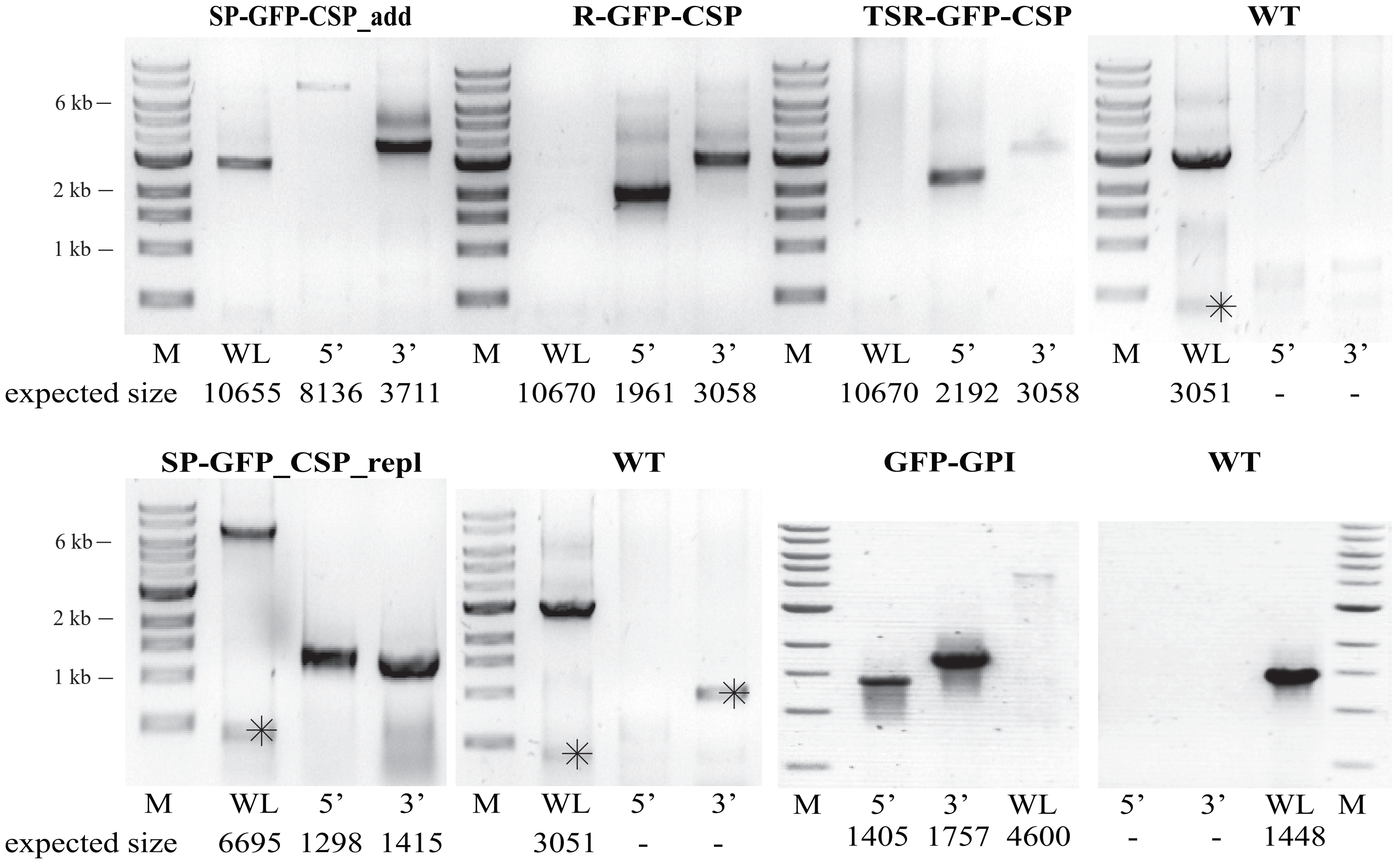

SP-GFP-CSP_add
R-GFP-CSP
TSR-GFP-CSP
WT
6 kb
2 kb
1 kb
M WL 5’ 3’ M WL 5’ 3’ M WL 5’ 3’ M WL 5’ 3’
expected size 10655 8136 3711 10670 1961 3058 10670 2192 3058 3051 - -
SP-GFP_CSP_repl
WT
GFP-GPI
WT
6 kb
2 kb
1 kb
M WL 5’ 3’ M WL 5’ 3’ M 5’ 3’ WL 5’ 3’ WL M
expected size 6695 1298 1415 3051 - - 1405 1757 4600 - - 1448

P99 CTAGCTAGCTTAATCATTCTTCTCATATACTT

P134 GAGCATACAAAAATACATGCACAC

P137 TGATTTACTTCCATCATTTTGCCC

P208 CCG**GAATTC**ATGTGTTGGTTGTAATTGAGG

P210 TTAA**CATCAC**CATCTAATTCAACAAG

P267 CCGGAATTCATGAGCACGCTTTTACTTTGTC

P268 GCAATTC**CATATG**TCCTCCTCCTCCTTGTCCATATCCTGGAAGTAGAGAATTAAC

P269 AAC**GGCGCC**GGTGGAGGTGGAGGTGGAGGTGGAAATAAAAGCATCCAAGCCCAAAG

P270 CCG**GATATC**CAGAAATATTTCAAAAGCCTACATAAC

P271 GCAATTC**CATATG**ACCTCCACCTCCACCTCCACCTCCACCACCTGGCTGTGGTTGTG

P272 AAC**GGCGCC**GGAGGTGGAGGTGGAGGTGGAGGTAATAACAATAACAAAAATAATAATAATGACG

P273 GCAATTC**CATATG**TCCACCTCCACCTCCACCTCCACCTGAACATTTATCCATTTTACAAATTTCAGTATC

P274 AAC**GGCGCC**GGAGGTGGAGGTGGAGGTGGAGGTAGTATATTTAATATTGTAAGCAATTCATTAGG

P882 AGGAGAATTAACCAATGCTGTATAC

P893 GCCGTCCTCGATGTTGTGGCGGATGACATATTACATGTTTGGGGATTTTTG
